# Assessment of Oxford Nanopore whole genome sequencing for large-scale genomic characterisation of *Staphylococcus aureus* bacteraemia isolates

**DOI:** 10.64898/2026.03.30.715209

**Authors:** Ingvild Haugan, Helene Marie Flatby, Hilde Lysvand, Nina Vibeche Skei, Kyriakos Zaragkoulias, Erik Solligård, Torunn Gresdal Rønning, Lene Christin Olsen, Jan Kristian Damås, Jan Egil Afset, Christina Gabrielsen Ås

## Abstract

Whole-genome sequencing (WGS) is increasingly being utilised in microbial diagnostics, surveillance, and research. In this paper we assess the performance of one leading long-read sequencing technology, Oxford Nanopore Technology (ONT), on 836 *Staphylococcus aureus* bacteraemia isolates. We compare the results to that of a leading short-read sequencing technology, Illumina.

All isolates were sequenced using ONT MinION Mk1B and Illumina HiSeq or MiSeq. Libraries were prepared according to manufacturers’ instructions. Preprocessing and downstream bioinformatic analyses were performed using a combination of in-house pipelines and publicly available software tools.

The average base substitution error rate in ONT assemblies was low but varied between sequence types, possibly due to lineage-specific methylation patterns. Multi locus sequence typing was similar between the technologies, while ONT assemblies allowed for better *spa* typing than Illumina assemblies. The reported detection rate was similar between ONT and Illumina assemblies for most virulence- and AMR-associated genes and variants. For 42 (22.2%) of 189 genes/variants, the two technologies disagreed in gene detection in 5 isolates or more, and in 39 (20.6.%) of these the highest detection rate was found with ONT. Discrepancies were mainly associated with low GC content, multiple repetitive segments, and small plasmids. Polishing of ONT data resulted in minor changes in gene/variant calling.

Our study supports the use of ONT WGS for bacterial population genomic studies on a large collection of *S. aureus* isolates. While assembly of ONT reads may be affected by its own methodological limitations, it was superior to Illumina assemblies in detection of potentially clinically relevant genes and variants at a low read error rate. Understanding the advantages and limitations of WGS technologies is essential before undertaking studies involving such methods on large sets of bacteria.

**Author summary:** In this paper, we present a practical assessment of one important whole genome sequencing (WGS) method, Oxford Nanopore Technology (ONT), and compare its performance in bacterial population genomics to that of WGS with Illumina technology. Our goal was to investigate the usefulness of ONT in studies aiming to identify clinically relevant bacterial characteristics in large collections of bacteria, such as genotype-phenotype studies. We sequenced a large set of clinical *S. aureus* isolates from episodes of bloodstream infections using both ONT and Illumina technologies and performed analyses with widely used software and bioinformatic pipelines.

We have elucidated inherent strengths and limitations of ONT and Illumina sequencing and report some of the practical consequences of these on bacterial typing and detection of clinically relevant genes. With this study, we present one of the most comprehensive assessments of long-read sequencing technology for the genomic characterisation of clinical bacterial isolates, and the findings provide guidance for researchers considering WGS in large-scale bacterial genomics.

## Introduction

Whole-genome sequencing (WGS) is increasingly being utilised in microbial diagnostics, surveillance, and research. While WGS may offer more detailed genomic information than any other diagnostic strategy, WGS technologies are often labour-intensive and expensive, and may require advanced equipment and highly specialised personnel. Before undertaking a project that involves WGS of large bacterial collections, it is important to know whether the chosen WGS technology is appropriate to answer one’s research question.

In the early 2000’s, sequencing technologies that allowed for high throughput and low turnaround time were developed and commercialised [1], expanding the accessibility of WGS to smaller research communities. These technologies, often referred to as next- or second-generation sequencing, usually require a polymerase chain reaction (PCR) in the sample preparation and/or when samples are immobilised onto a flow cell [2]. Most second-generation sequencers identify nucleotides as complementary strands to DNA fragments as they are built and achieve high output through massive parallel sequencing.

The second-generation sequencing technology Illumina uses a sequencing-by-synthesis method with bridge amplification and clustering of identical molecules [1]. This results in a strong signal for each fluorescently labelled nucleotide and a low base substitution error rate with base calling accuracy exceeding 99.99% [3, 4].

High accuracy is a main advantage of this technology, but the error rate remains low only for a limited number of cycles as the clusters eventually lose synchrony [5]. To ensure high base quality in all reads the read lengths are therefore preset and relatively short, typically 150– 300 base pairs [4, 6]. Some more recently developed Illumina platforms offer high throughput at the cost of read length to maintain a low error rate. While there are some differences in the base substitution error rate between them, the accuracy remains high [7]. The most recent Illumina platform, the MiSeq i100 Series, provides longer read lengths (2×500 pb) with lower output, but its accuracy has not been assessed in a peer reviewed article.

Due to the short lengths of the reads, complete genomes cannot be assembled from second-generation sequencing [8]. Larger genomic variations, such as inversions and translocations, as well as repetitive regions, are particularly challenging to assemble correctly from short reads.

Technologies that sequence without DNA fragmentation and PCR were introduced from 2009 and onwards, and are often referred to as third-generation, or single molecule sequencing (SMS) [1, 2]. One major advantage of such platforms is the generation of very long reads, ranging from several thousand to a couple of million base pairs (bp), which allows them to overcome the difficulties of short-read sequencing with structural variants and low complexity regions and thus enables the assembly of complete genomes [8].

With Oxford Nanopore Technology (ONT) sequencing, the DNA strand is read as it travels through a protein nanopore, disrupting the ionic current across a membrane [1]. This technology can offer ultralong read lengths, and the first read exceeding 2 Mbp was published in 2018 [9].

Long-read sequencers, including ONT, have so far not achieved the base calling accuracy of short-read sequencing. One strategy to circumvent higher read error rates is to polish long-read assemblies with short-read data [10], an expensive and time-consuming strategy, particularly when working with a large collection of bacteria. The technologies are, however, evolving rapidly, and substantial progress have been made in recent years. The introduction of R10 flow cells, new base calling models, and duplex base calling are notable advances in ONT technology that have increased the accuracy to levels that may render polishing with short-read data obsolete [11–13].

In this study we assess the performance of a leading long-read sequencing technology, ONT, on genomic characterisation of a large collection of clinical *Staphylococcus aureus* isolates and compare it to the results from the more widely used short-read sequencing technology, Illumina. Both these technologies are increasingly being used in large-scale comparative genomic studies of bacteria, including genome wide association studies (GWAS), and it is thus essential to understand the practical implications of their inherent strengths and weaknesses when selecting a sequencing platform.

Rather than conducting a benchmarking study, we concentrate on comparing quality metrics and results from genotyping and gene calling using a set of widely used assembly, polishing, and bioinformatic tools. We have particularly focused on, and further explored discrepancies in, detection of genes that may be relevant for antimicrobial resistance and virulence. This is, to our knowledge, the first study to perform large-scale genomic comparison of over 800 clinical bacterial isolates sequenced by ONT.

## Material and methods

### Strain collection

The Nord-Trøndelag Hospital Trust Sepsis Registry has included all episodes of bacteraemia in patients admitted to any of the hospitals in the region, Levanger and Namsos hospitals, since 1994. A total of 923 unique *S. aureus* bacteraemia (SAB) episodes in patients 18 years or older were registered between 1996-2022, and we retrieved bacterial isolates from 860 of those, out of which 836 were sequenced with both Oxford Nanopore and Illumina technologies and included in this study.

### Whole genome sequencing and assembly

Briefly, strains were grown overnight on sheep blood agar plates at 35 °C. Cells were treated with proteinase K (2 mg/mL) and lysostaphin (0.1 mg/mL) for 15 min with shaking at 37°C, before heating for 15 min at 65°C. Genomic DNA was then isolated using the EZ1 DNA tissue kit with an EZ1 Advanced XL instrument (Qiagen).

Long-read sequencing libraries were prepared and multiplexed using the Rapid Barcoding kit V14 (SQK-RBK 114) according to the manufacturer’s instructions. Sequencing libraries were sequenced with R10 flow cells (FLO-MIN114) on a MinION Mk1B sequencer (Oxford Nanopore Technologies). Base calling was performed using Dorado v0.7.2 with the dna_r10.4.1_e8.2_400bps_hac@v5.0.0 base calling model. Base called reads from ONT were quality controlled with NanoComp v1.24.0, filtered using filtlong v0.2.1 (--min_length 1000 --keep_percent 95) (https://github.com/rwick/Filtlong) and assembled with Flye v2.9.4 (--nano-hq) [14].

Short-read sequencing libraries were prepared using Nextera XT according to the manufacturer’s instructions and were sequenced on an Illumina HiSeq or NovaSeq sequencer (2×150 bp). FastQC [15] was used for quality control, Trimmomatic for trimming/filtering (illuminaclip, leading:10 trailing:10 minlen:30) (https://github.com/usadellab/Trimmomatic) and Shovill [16] for assembly using the Nullarbor v2 pipeline.

Nanopore assemblies were additionally polished with Illumina data using Pypolca v0.3.1 (--careful) [10, 17]. All assemblies were quality checked using QUAST v5.2.0 (https://github.com/ablab/quast).

A nanopore assembly was interpreted as a complete chromosome if the genome size was >2 650 000 bp and was reported as circular by the assembler.

### Bioinformatic analyses

For annotation, *spa* typing and sequence typing, we used Prokka [18], spaTyper (https://doi.org/10.5281/zenodo.4063625) and mlst [19], with mlst assigning sequence types using curated PubMLST schemes [20]. For calling of AMR- and virulence genes as well as plasmid replicons, we used AMRFinderPlus (AMRF+) [21], Abricate [22] with the Virulence Finder Database (VFDB) [23] and PlasmidFinder database [24]. For any genes or variants present in both AMRF+ and VFDB we only included results from AMRF+. We refer to gene/variant copy number as the number of gene/variant copies reported in each isolate. The gene/variant presence is reported as the detection of a gene or variant in one isolate, ignoring multiple copies. Detailed information about reported genes and variants is listed in Supplementary Table S1.

Raw data from isolates with gene/variant discrepancies between ONT and Illumina were mapped against the respective reference gene and coverage assessed using minimap2/bwa [25] and samtools [26, 27] accordingly.

High error sequence contexts were identified by aligning filtered ONT reads to their respective error-corrected reference genomes using minimap2 v2.28, processing alignments with samtools v1.18, calling variants with bcftools mpileup and bcftools call, filtering high-confidence SNPs with bcftools filter [28], and extracting error-enriched regions using bedtools v2.31.1 [29]. For motif discovery, sequence contexts of high-error isolates (≥25 base errors/1 000 000 bases) were compared with those of low-error isolates (≤7 base errors/1 000 000 bases) using DREME [30] and STREME [31] (MEME Suite v5.5.2) with an E-value threshold of 0.1.

## Results

### Quality

The mean average ONT read length and quality score were 6 042 base pairs (bp) and 16.3, respectively (Table 1). The mean GC content of ONT reads was on average 32.9% compared to 36.1% in Illumina reads. The mean genome size of ONT assemblies was 2 787 678 bp which was 2% larger than for Illumina assemblies. A complete chromosome was assembled from ONT reads in 807 (96.5%) isolates. In 20 (2.4%) ONT assemblies the largest contig was >2 650 000 bp but not circular. The median number of contigs was 2, with four or fewer contigs in 827 (99.0%) isolates. Two outlier isolates had >54 contigs, had the lowest mean coverage of 8.9 and 10.6 respectively, and were the only isolates with N50 < 1 000 000.

**Table 1.** Quality of ONT and Illumina raw data and assemblies.

|  | ONT<br>Mean (range) | Illumina<br>Mean (range) |
| --- | --- | --- |
| **Raw data** |  |  |
| Average read length | 6 042 (2 729-13 314) | 150 |
| GC% | 32.9 (32.7-33.0) | 36.1 (34.0-39.1) |
| Coverage | 137 (9-570) | 461 (118-1 100) |
| Average quality score | 16.3 (13.0-19.0) | 37.6 (31.8-39.6) |
| **Assemblies** |  |  |
| Number of contigs | 1.8 (1-84), median: 2 | 123 (12-488), median: 103 |
| Genome size | 2 787 678 (2 650 808-3 025 323) | 2 733 455 (2 627 570-2 870 907) |
| N50 | 2 758 527 (59 870-2 950 353) | 80 653 (10 678-671 975) |

### *In silico* genotyping

A total of 63 different sequence types (ST) were identified in 747 (89.4%) isolates after polishing ONT assemblies (Figure 1). Of these, 740 (88.5%) were identified from ONT and 742 (88.8%) from Illumina assemblies. One (0.001%) isolate successfully sequence typed from Illumina assemblies was not typed from neither ONT nor polished assemblies due to the presence of a duplicated allele. In 18 (2.2%) of the remaining 88 (10.5%) isolates the locus combinations did not correspond to a known ST. The last 70 (8.4%) isolates could not be sequence typed due to ambiguous alleles.

**Figure 1.**
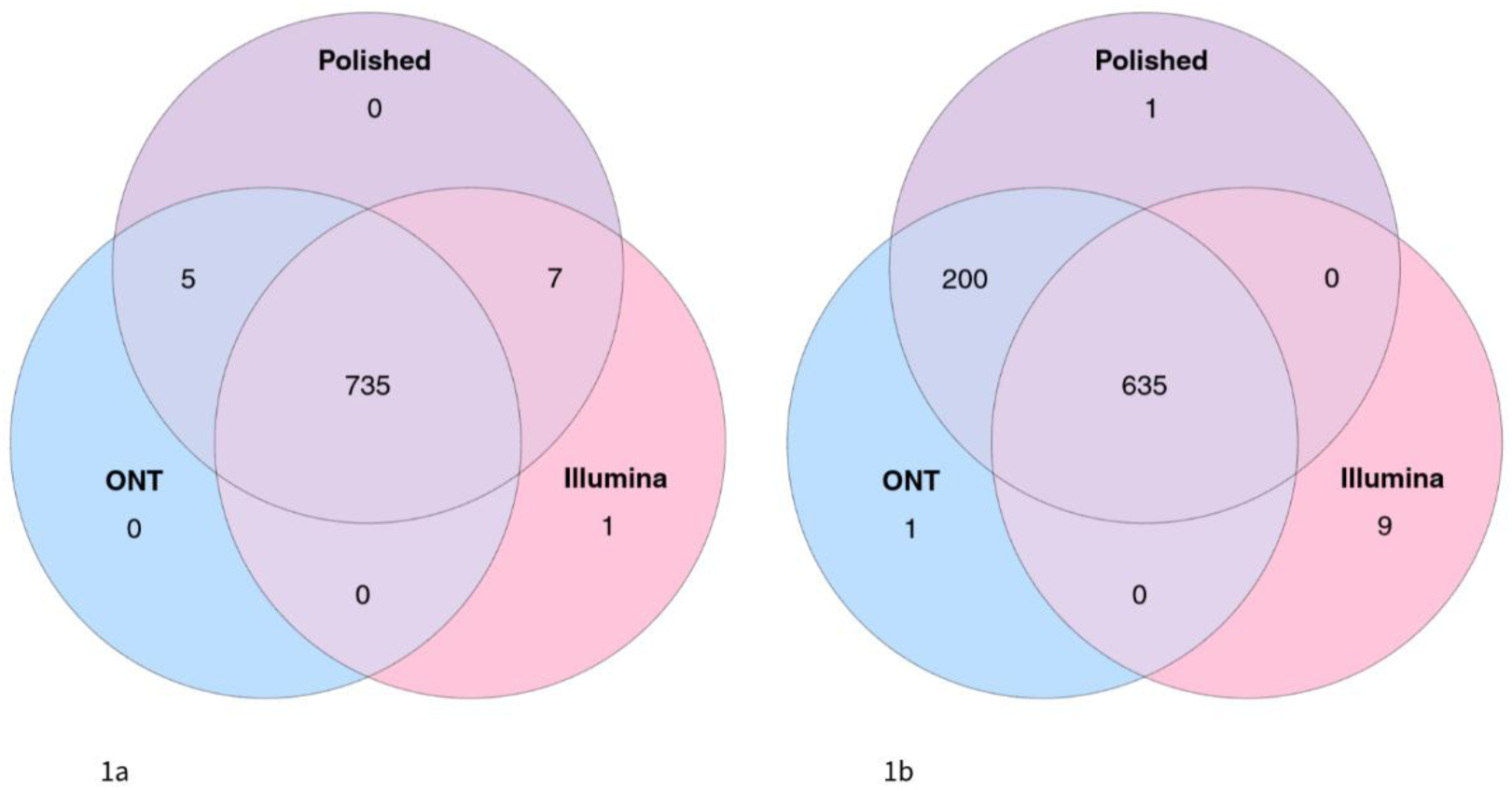
Number of isolates with concordant and discordant ST (1a) and *spa* types (1b) between ONT, Illumina and polished assemblies

In polished assemblies, all isolates were assigned to a *spa* type (n=797, 95.3%) or a novel *spa* repeat sequence (n=39, 4.7%) (Figure 1). Results from ONT assemblies correlated with polished results in 835 (99.9%) isolates and reported a different spa type in one (0.001%) isolate. From Illumina assemblies, a *spa* type or *spa* sequence concordant with the polished results was found in 636 (76.1%) isolates. In the remaining 192 (23.0%) isolates a *spa* type could not be confidently identified as the *spa* gene was fragmented on multiple contigs. Illumina reported another eight *spa* types and one *spa* repeat that were not in concordance with the results from polished ONT. In all of these, the *spa* type in the polished ONT assemblies consisted of a high number of *spa* repeats, of which some had been collapsed and misassembled from Illumina reads. In addition, three of these *spa* repeats also had nucleotide differences resulting in one or two discordant repeat types in the *spa* genes from Illumina assemblies compared to ONT.

### Base errors and indels in ONT assemblies

After polishing ONT reads with Illumina data, the median number of corrected base and insertion/deletion (indels) errors per 1 000 000 bases were 2.6 and 3.6, respectively (Table 2).

**Table 2.**
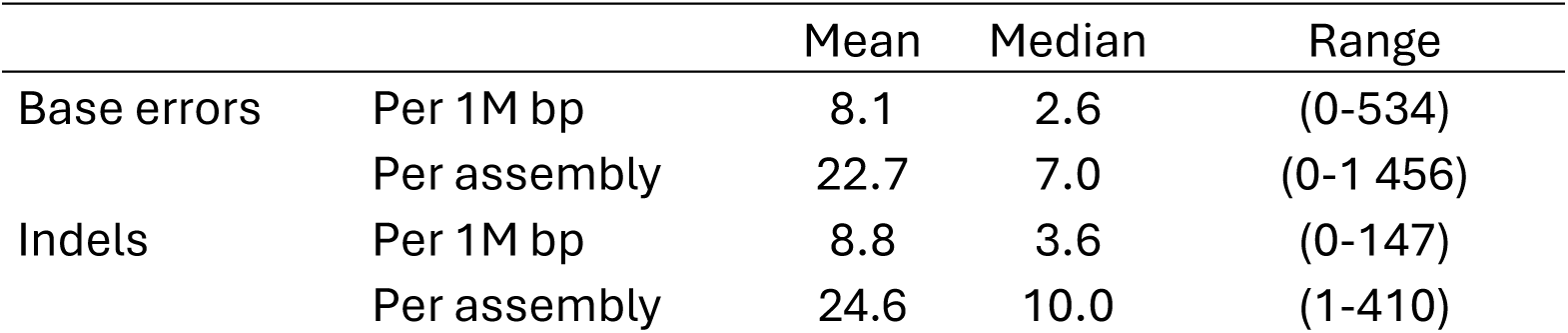
Errors per 1 000 000 bp and per assembly in Oxford Nanopore assemblies.

|  |  | Mean | Median | Range |
| --- | --- | --- | --- | --- |
| Base errors | Per 1M bp | 8.1 | 2.6 | (0-534) |
|  | Per assembly | 22.7 | 7.0 | (0-1 456) |
| Indels | Per 1M bp | 8.8 | 3.6 | (0-147) |
|  | Per assembly | 24.6 | 10.0 | (1-410) |

The highest error rates were observed in isolates belonging to ST25, which had mean base and indel error rates of 51.7 (36-73) (Figure 2) and 104.5 (87-147) per 1 000 000 bases, respectively.

**Figure 2.**
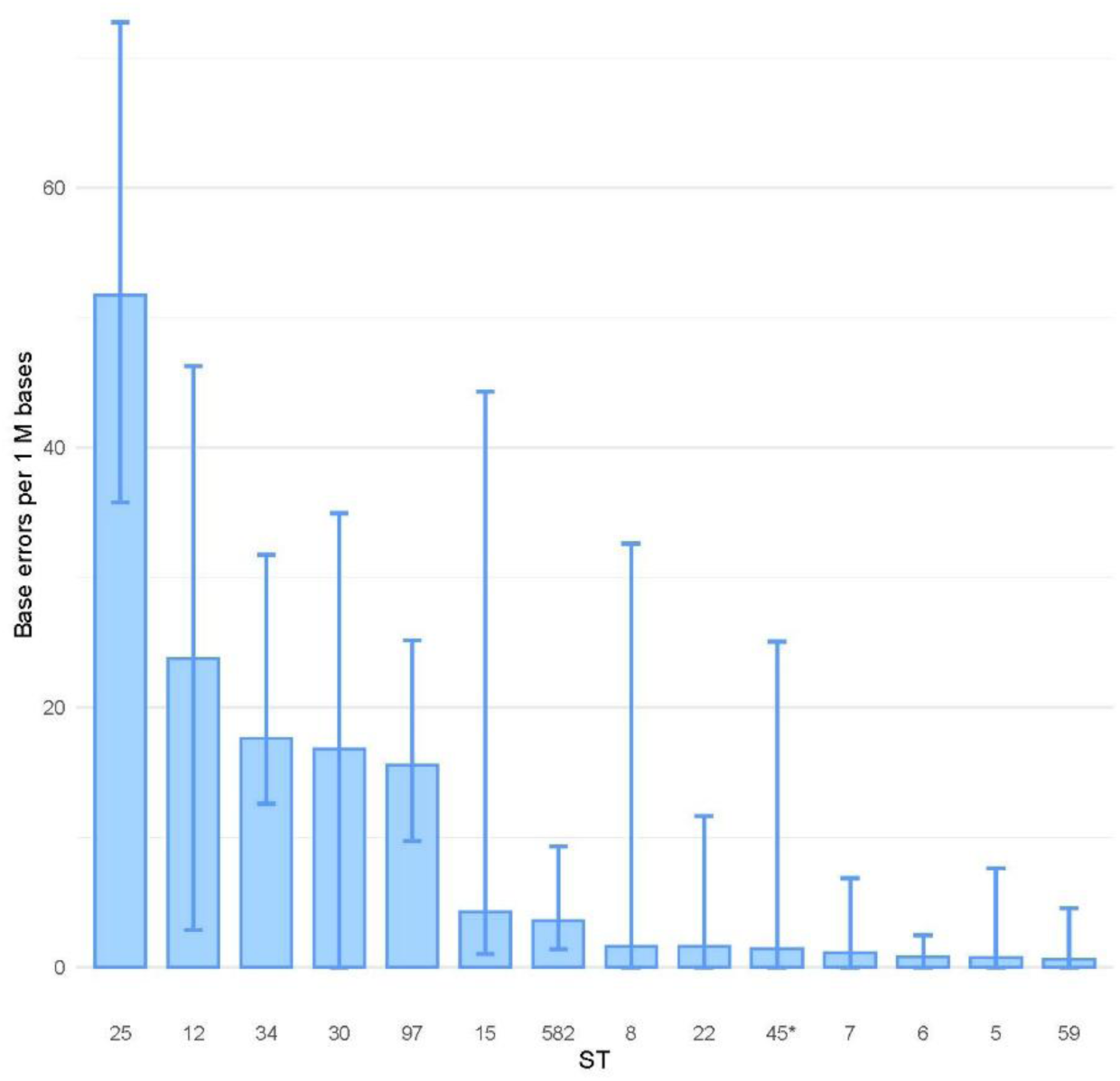
The mean number and range of base errors per 1 000 000 bases in isolates from sequence types (ST) with ≥10 isolates in total *Excluding one outlier isolate from ST45 with 534 base errors and 71 indels per 1 000 000 bases

In total, 38 (4.6%) isolates were categorised as isolates with high base error rate and all isolates in ST25 (n= 24) were in this group. The remaining 14 high error isolates belonged to six other specific STs or non-typable ST. Sequence type 25 did not otherwise deviate from other STs in quality markers (Table 1 and Supplementary Table S2). Motif enrichment analysis using STREME detected 17 motifs significantly associated with high base error rates (Supplementary Figure F1). The e-value of the top motif, DWGGWCCWH, was 9.3×10^-70^ (p <0.001).

### Calling of antimicrobial resistance and virulence genes

Of all antimicrobial resistance (AMR), virulence, and stress genes and variants identified in polished ONT assemblies (total gene count, n=81 309), 81 306 (99.99%) and 78 836 (96.96%) were detected in ONT and Illumina assemblies, respectively. Ignoring multiple copies of genes/variants in some isolates, 75 380 (100.0%) of the gene/variant presence detected in polished assemblies (gene presence, n=75 383) was identified in ONT assemblies compared to 74 134 (98.3%) in Illumina assemblies (Figure 3).

**Figure 3.**
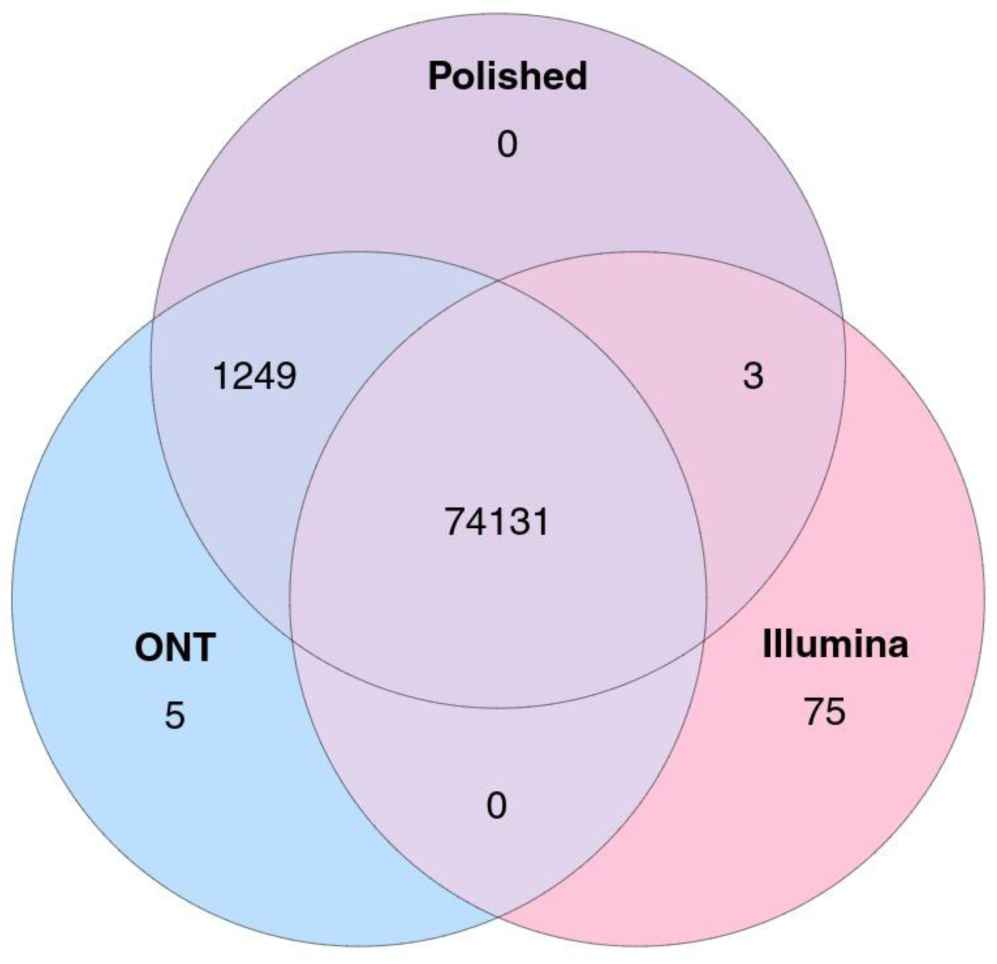
Concordance and discordance in reported presence of genes/variants in isolates between ONT, Illumina and polished assemblies

A total of 189 different AMR, virulence, and stress genes/variants were reported, 111 (58.7%) from the AMRF+ database and 78 (41.3%) from VFDB (Supplementary Table S1). On average, a total of 97.3 genes were reported per isolate in ONT compared to 94.4 in Illumina assemblies. For 99 (52.4%) of the genes the presence/absence and copy number were consistent between ONT, Illumina, and polished ONT assemblies.

For 82 (43.4%) of the 90 (47.6%) genes/variants with discrepancies in presence/absence and/or copy number between ONT and Illumina, the reported gene copy number was higher from ONT assemblies. Compared to polished ONT assemblies, the presence of a gene/variant was detected in ONT and missed in Illumina assemblies for 68 (36.0%) genes/variants in 1-120 isolates. For each of the genes *arsB, cna* and *sea*, one ONT assembly failed to detect the gene despite its presence in the respective Illumina ad polished assemblies. The genes with discrepant detection in five isolates ore more are depicted in Figure 4.

**Figure 4.**
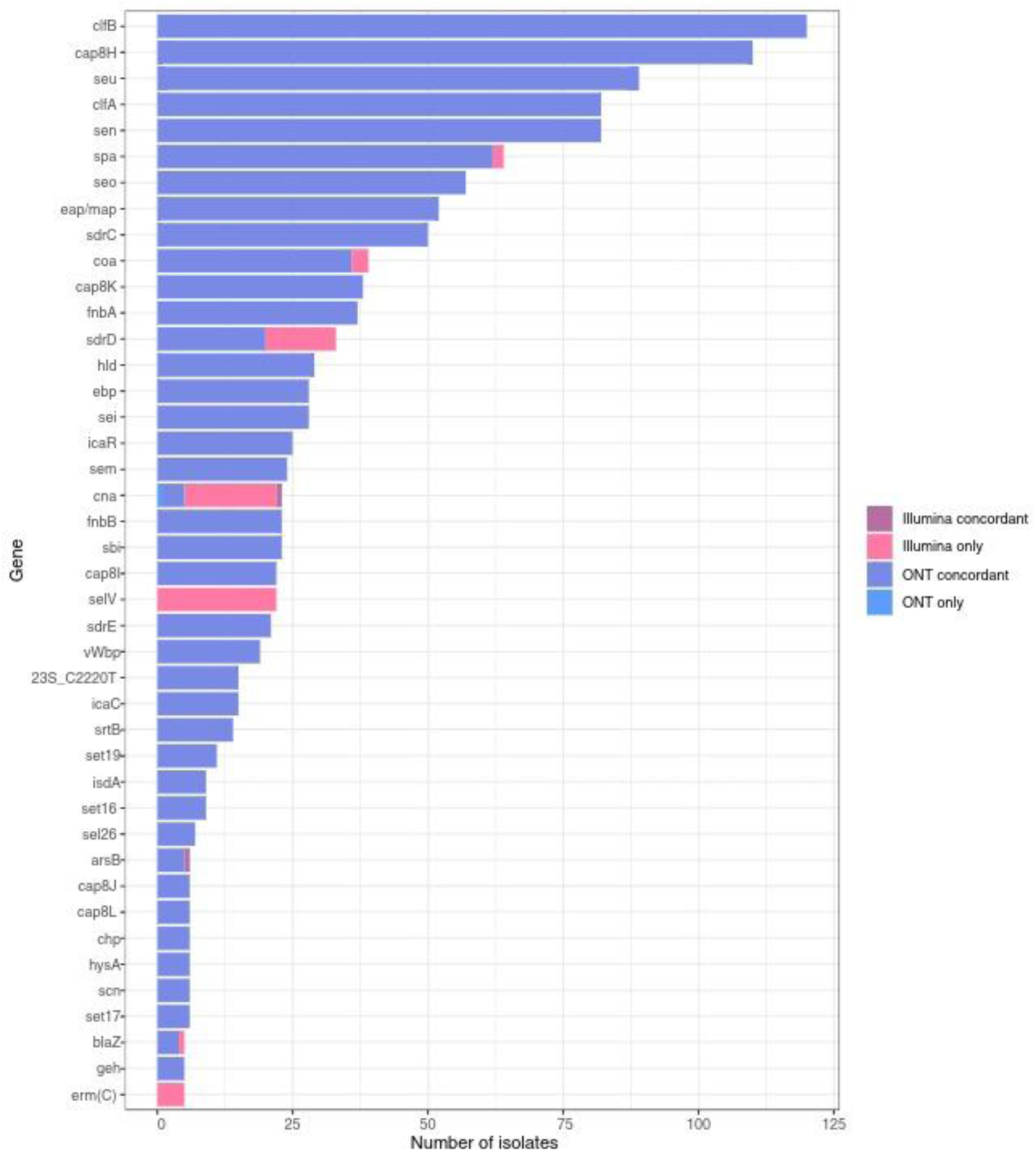
Genes with highest discrepancies in detection rate between assemblies (≥5 isolates), showing the number of isolates with gene/variant detection in only one technology, either concordant (blue: ONT, mauve: Illumina) or discordant (medium blue: ONT, pink: Illumina) with polished assemblies.

Discrepancies in gene detection between the technologies were higher for virulence and stress genes than for AMR genes (Table 3). Seventeen (9.0%) genes/variants that were not present in the polished ONT assemblies were called in the Illumina assemblies of 1-22 isolates. In comparison, two (1.1%) such genes/variants were reported in the ONT assemblies of 1-4 isolates. For 12 (6.4%) of the genes/variants with no discrepancy in the gene presence between technologies, the gene count was lower in Illumina assemblies than in ONT and polished assemblies due to multiple gene copies. The genes/variants with highest discrepancies in reported presence (in absolute numbers) and gene count between ONT and Illumina are listed in Table 3. For a complete list of reported detection rate and copy count of genes, discrepancies, and comparisons to polished ONT, mapping, and reference genes, see Supplementary Table S1.

**Table 3.** Number and proportion of isolates with highest discrepancies in reported genes/variants and/or total gene/variant count for each gene/variant category, corrected results*, and GC content of reference genes.

|  | ONT |  | Illumina |  | Corrected* |  | Ref. gene |
| --- | --- | --- | --- | --- | --- | --- | --- |
|  | Presence<br>n (%) | Copy no<br>n | Presence<br>n (%) | Copy no<br>n | Presence<br>n (%) | Copy no<br>n | GC content<br>% |
| **AMRF+ amr genes** |  |  |  |  |  |  |  |
| 23S_C2220T | 15 (1.8) | 28 | 0 (0.0) | 0 | 15 (1.8)^ | 28^ | 49.9 |
| *erm(C)* | 7 (0.1) | 10 | 12 (1.4) | 13 | 12 (1.4)^ | 16^ | 25.6 |
| ABC-F | 313 (37.4) | 313 | 309 (37.0) | 309 | 309 (37.0) | 309 | NA<sup>§</sup> |
| *blaZ* | 581 (69.5) | 714 | 578 (69.1) | 577 | 581 (69.5) | 714 | 27.9 |
| **AMRF+ virulence/stress genes** |  |  |  |  |  |  |  |
| *seu* | 510 (61.0) | 510 | 421 (50.4) | 421 | 510 (61.0) | 510 | 26.8 |
| *sen* | 501 (59.9) | 501 | 419 (50.1) | 419 | 501 (59.9) | 501 | 25.4 |
| *seo* | 364 (43.5) | 364 | 307 (36.7) | 307 | 364 (43.5) | 364 | 24.5 |
| *hld* | 834 (99.8) | 834 | 805 (96.3) | 805 | 834 (99.8) | 834 | 30.9 |
| **VFDB virulence/stress genes** |  |  |  |  |  |  |  |
| *clfB* | 820 (98.1) | 820 | 700 (83.7) | 700 | 820 (98.1) | 820 | 39.2 |
| *cap8H* | 647 (77.4) | 647 | 537 (64.2) | 537 | 647 (77.4) | 647 | 27.2 |
| *clfA* | 831 (99.4) | 831 | 749 (89.6) | 749 | 831 (99.4) | 831 | 39.3 |
| *spa* | 597 (71.4) | 597 | 537 (64.2) | 537 | 597 (71.4) | 597 | 38.1 |
| *eap/map* | 824 (98.6) | 824 | 772 (92.3) | 772 | 824 (98.6) | 824 | 27.5 |
| *sdrC* | 818 (97.9) | 818 | 768 (91.9) | 768 | 818 (97.9) | 818 | 35.3 |
| *cap8K* | 650 (77.8) | 650 | 612 (73.2) | 612 | 650 (77.8) | 650 | 26.9 |
| *fnbA* | 685 (81.9) | 685 | 648 (77.5) | 648 | 685 (81.9) | 648 | 37.1 |
| *esaG* | 836 (100.0) | 5 868 | 836 (100.0) | 5 036 | 836 (100.0) | 5 868 | 29.7 |

\*Corrected against polished ONT and/or mapping, ^mapped, <sup>§</sup>Not Applicable

Among the 53 AMR genes/variants, there were discrepancies in reported presence and/or gene count between ONT and Illumina assemblies in 12 (22.6%). The discrepancy was highest for the oxazolidinone resistance associated 23S rDNA C2220T variant, the macrolide-lincosamide-streptogramin B (MLS) resistance gene *erm(C)*, the beta-lactam resistance gene *blaZ*, and an ATP-Binding Cassette-Family (ABC-F) protein encoding gene (Table 3).

The presence of a 23S rDNA C2220T variant was detected in 15 (1.8%) isolates from ONT assemblies and was absent in Illumina assemblies. Mapping of Illumina data to the 23S rRNA gene revealed that the C2220T mutation was present in 31.6-35.2% of reads (coverage 411-2 322), corresponding to 2 copies each in the 15 discrepant isolates. In ONT assemblies, the mutation was detected in 2 copies of the 23S rRNA gene in 13 isolates and in a single copy in 2 isolates. More information on results from mapping of these 15, and other selected, discrepant isolates is found in Supplementary Table S3.

The *ermC* gene was detected in 12 (6.4%) and 7 (3.7%) isolates from Illumina and ONT assemblies, respectively. For the 5 discrepant strains, mapping of ONT reads to the *ermC* gene confirmed its presence, all with an average gene coverage >6 times the average genome coverage, consistent with a plasmid location of the gene (Supplementary Table S3).

The *blaZ* gene was called from 581 (69.5%) ONT assemblies compared to 578 (69.1%) Illumina assemblies. One of the *blaZ* genes found with Illumina was not present in ONT assemblies. Mapping of ONT (n=1) and Illumina (n=4) raw reads to the *blaZ* gene confirmed its presence in all five isolates with gene coverages <19% of the average genome coverage in the Illumina data, and >300% of the average genome coverage in the ONT data (Supplementary Table S3). In the isolate missing *blaZ* in the ONT assembly, the replicon repUS5_1_CDS20(pETB) was found on the same contig as the *blaZ* gene in the Illumina assembly, suggestive of a plasmid localisation (Supplementary Table S4).

An ABC-F protein family gene was reported in 313 (37.4%) of ONT assemblies and 309 (37.0%) of Illumina assemblies. The reference gene in the AMRF+ database was *optrA*, a gene that is rarely found in *S. aureus* [32]. Given that all hits had very low identity scores of 35-36% to the reference gene, we have disregarded this result.

In 78 (57.4%) of the136 virulence and stress genes reported, we observed discrepancy in reported presence and/or gene count between technologies. Discrepancies in presence were often observed for enterotoxin genes, capsular genes, and adherence genes (Table 3). The gene/variant with the highest gene count discrepancy was the antitoxin gene *esaG*.

The adherence genes *clfA* and *clfB* were reported in 831 (99.4%) and 820 (98.1%) ONT assemblies, respectively (Table 3). The corresponding results in Illumina assemblies were 749 (89.6%) and 700 (83.7%). Mapping of Illumina reads from 16 randomly selected isolates lacking *clfA* and/or *clfB* to the respective genes confirmed their presence with average gene coverages between 2.1 and 5.2 times the average genome coverages. Similar results were found when mapping Illumina reads of selected isolates to the adherence genes *sdrC* and *fnbA* (Supplementary Table S3).

The capsular gene *cap8H* was found in 647 (77.4%) isolates from ONT assemblies and 537 (64.2%) isolates from Illumina assemblies. Mapping of Illumina reads from five of the discrepant isolates to the reference gene revealed its presence, however with coverages <17% of the average genome coverage in all isolates.

Seven of the ten AMRF+ virulence/stress genes with the highest discrepancy in presence between ONT and Illumina assemblies were staphylococcal enterotoxin (SE) genes. For 16 of the 29 genes coding for enterotoxins and enterotoxin-like proteins, we observed discrepancies in reported detection rate between the technologies (Table 4). The mean SE reference gene GC content was 28.0% (24.5%-31.1%) in those with discrepant results and 30.9% (26.1%-45.8%) in those with equal results (Table 4, Supplementary Table S1). While both technologies detected 1-12 types of SE genes in the assemblies of the same 686 (82.1%) isolates, the types of enterotoxin genes differed in 221 (26.4%) of these.

**Table 4.**
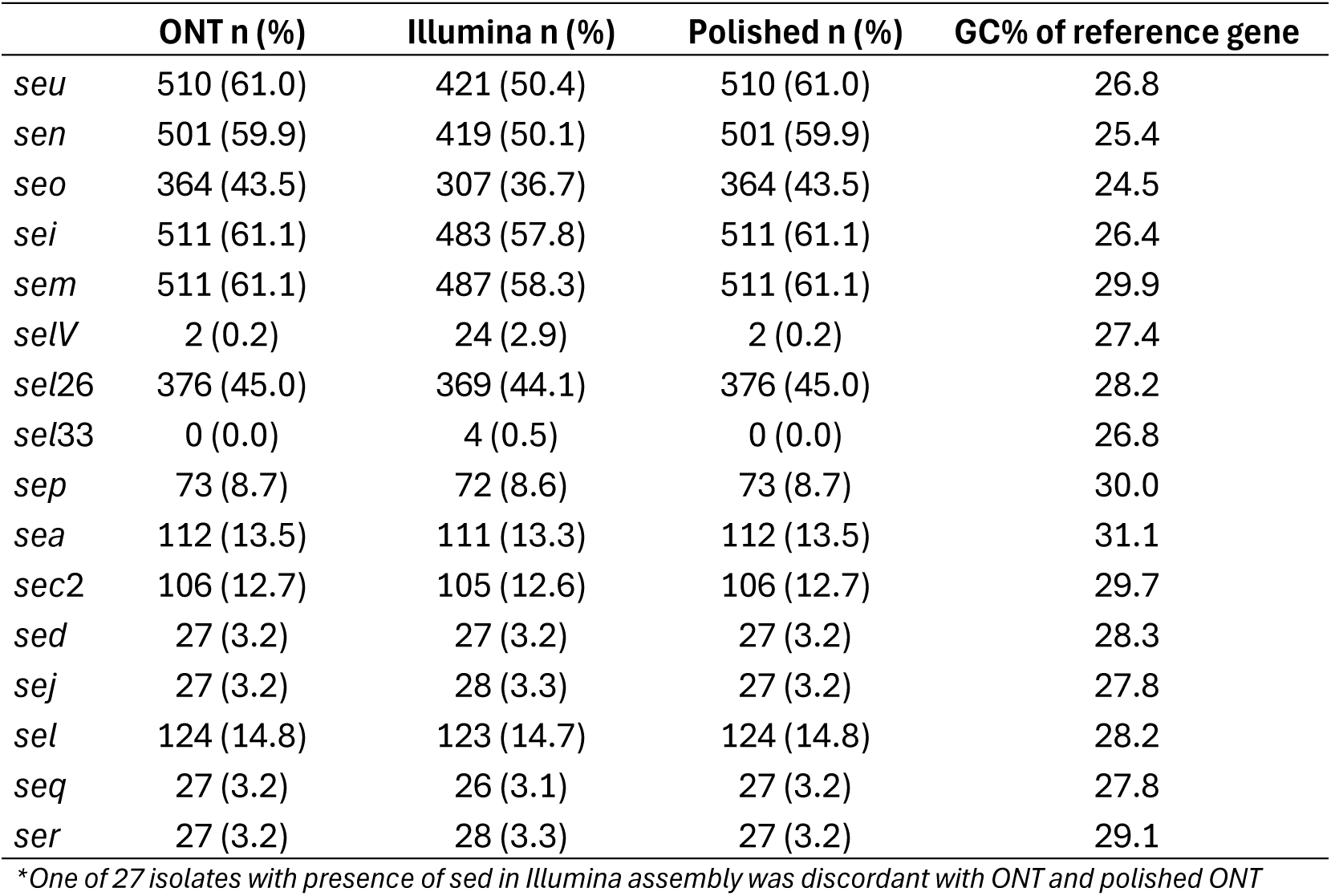
Number and proportion of isolates with discrepancies in detection of staphylococcal enterotoxin genes between ONT and Illumina, compared to polished ONT assemblies, and GC content of reference gene.

In 10 randomly selected isolates where the *seu* gene was detected only in ONT assemblies, mapping of Illumina data to the reference gene revealed its presence with a gene coverage of less than 9% of the genome coverage in all isolates. Mapping of ONT raw reads of 10 isolates from which the *selV* gene was only detected in Illumina assemblies confirmed the presence of this gene with gene coverages similar to that of the genome in all isolates.

## Discussion

We have presented data supporting an overall good performance of ONT in genotyping and AMR and virulence gene calling of 836 clinical *S. aureus* isolates. Our results show that high quality assemblies enabling reconstruction of complete, or near-complete, bacterial genomes, can be achieved from ONT sequencing. This offers opportunities for analysing the genomic context, including organisation and mobility of genetic elements, which may be important for antibiotic resistance and virulence. However, there were other notable discrepancies between results from ONT and Illumina that can have considerable implications for comparative genomic studies.

Previously published results, mainly on smaller bacterial collections, have highlighted the advantages of combining ONT and Illumina in bacterial genome sequencing [4, 8, 10], however this strategy may not be feasible for larger studies. In this paper we have assessed the utility of ONT for studies that aim to perform comparative genomic analyses and GWAS on large bacterial collections. To address this, we have quantified ONT base calling errors and compared ONT genotyping and gene calling of possibly clinically relevant genes in a large set of *S. aureus* isolates with results from Illumina and polished ONT.

Previous studies have found an ONT base calling accuracy high enough for outbreak investigations when using the latest ONT chemistry combined with advanced bioinformatic tools [11, 33]. We detected no significant difference in sequence typing between the two technologies, whereas several isolates received a *spa* type from ONT assemblies and not from Illumina assemblies.

The rate of base correction when polishing ONT assemblies in our study was low overall with average base calling and indel rates consistent with a read accuracy beyond 99.9%. We observed a variation in error rates between isolates, and assemblies with high error rates did not seem to be randomly distributed among the different STs. This finding suggests that there are clone specific characteristics that may impact on the quality of ONT sequencing. Previous reports support variation in ONT performance between different bacterial species and Dabernig-Heinz et al (2024) also found different intraspecies error rates in 18 *S. aureus* isolates [4, 34, 35]. As far as we know, there are no other reports on ONT performance variation between subgroups within one species.

It has previously been shown that methylated nucleotides caused by restriction-modification systems in bacteria can cause ambiguity in ONT base calling [35] and this is one possible explanation for poorer base calling accuracy. Indeed, our enrichment analyses revealed a significant association between several motifs and high-error regions. The top hit was a motif recognised by the type II methyltransferase M.Sau961 [36] supporting methylation as a likely explanation for high error rates in some subgroups of isolates. Methylation motifs differ between different *S. aureus* lineages [37], and to our knowledge, methylation motifs for ST25 have not been previously published. Further investigations are required to confirm whether the remaining motifs found in our study are associated with methylation.

If base errors occur within the reading frames of genes, they may interfere with gene calling. In our study, we detected more genes in ONT assemblies than in Illumina assemblies and polishing of ONT assemblies only corrected the gene count in a few cases. This suggests that the base read error is not a major driver of missing genes in ONT assemblies, but rather that mechanisms related to the assembly or annotation may explain these discrepancies.

Repetitive regions is a well-known problem in assembly of short-read sequences, particularly when they extend beyond the read lengths [38–40]. Like repetitive segments, reads from multicopy genes can be collapsed in the assembly of short reads, resulting in an underestimated gene count [41]. Our results demonstrate that long-read sequencing outperforms short-read sequencing in resolving such repetitive regions, as we observed high discrepancies in genes associated with multiple repeat regions and multicopy genes, such as the *clfA*, *clfB*, and *esaG* genes. This also explains why *spa* typing is often unsuccessful from Illumina assemblies.

High discrepancy genes not associated with repetitive regions, including some capsular polysaccharide synthesis (*cap*) and SE genes, were frequently characterised by low GC content. As Illumina sequencing is known to exhibit GC-dependent coverage bias [42, 43], this likely contributed to the low gene coverage in our Illumina data and the subsequent failure to assemble these regions.

The majority of the genes with high discrepancy were more often detected by ONT than Illumina, but the SE gene *selV* makes one exception, which cannot be explained by extreme GC content. The group of enterotoxin and enterotoxin-like protein encoding genes consists of several genes, some of which have been described only in recent years [44] and may be found in clusters of multiple genes in one isolate [45]. The products of SE genes share common phylogenetic relationships and have been allocated to groups based on the primary structure of the protein, a grouping that has evolved with the identification of additional toxins [44, 46]. The toxins share several conserved regions and typically consist of a conserved junction between two more variable domains that give the emetic and superantigen effects, respectively [46]. Further investigations are required to assess whether high degree of genetic similarity between some SE genes interfere with the assembly or annotation of such genes in ONT sequencing.

We found less disagreement between ONT and Illumina for AMR genes than virulence and stress genes. The 23S rDNA C2220T variant was probably discarded as a base read error in Illumina assemblies because the mutation occurred in only 1-2 of the 5-6 copies of this gene in each isolate that harboured the variant. The loss of small plasmids in assembly of long reads is a known problem [47] and can explain why the *ermC* gene was missed in some of our ONT assemblies, as this gene is often located on small plasmid.

## Limitations

This paper presents analyses of results from ONT and Illumina sequencing as they were performed in our laboratory, using the kits and instruments that were standard at the time, and a combination of in-house and published bioinformatic pipelines. However, sequencing technologies continue to evolve rapidly, including advances in library preparation chemistries, base calling algorithms, and assembly methods. These developments have the potential to substantially improve read accuracy, coverage uniformity, and downstream assembly quality. On the other hand, one advantage of choosing a more practical approach is that it may increase the generalisability to other research projects involving large sets of bacterial isolates that are also subject to financial, resource, and time limitations.

Our ONT and Illumina results were compared to ONT assemblies polished with Illumina data to identify base errors and gene calling discrepancies. Generating a consensus assembly by integrating results from multiple assemblers and combining both long- and short-read data remains a resource-intensive and computationally demanding process but might be regarded as the gold standard [48, 49]. It is possible that we would have exposed a higher degree of discrepancy between the technologies in a comparison against a true hybrid assembly.

Although polishing has the potential to introduce some base-level errors, this effect is likely limited if the Illumina data has sufficient depth (≥25x) [10]. While the overall Illumina read depth were substantially higher than 25x in all our isolates, we observed very low coverage in some extreme GC regions, and such regions may still introduce errors. The polisher used in our study (Pypolca-careful) was, however, found to rarely decrease overall accuracy.

In addition, the quality of Illumina assemblies would likely have improved if we had used the newer Illumina DNA prep kit rather than Nextera XT for library preparation, as the latter has been documented to introduce more GC-dependent bias leading to uneven sequencing coverage [50, 51] and thus affecting genome assembly.

As our collection of bacterial isolates was large and the genetic analyses extensive, in-depth analyses of mechanisms behind all gene discrepancies were not feasible. Though beyond the scope of this article, we believe investigations should be made to better understand such systematic errors and the consequences they may have for large-scale genomic studies of bacterial pathogens.

## Conclusions

Our study supports the use of ONT sequencing in genomic studies of larger sets of bacterial isolates and suggests that polishing with Illumina data only results in minor improvements in gene calling and has little effect on genotyping of *S. aureus*. Base calling errors may however not occur randomly, and sequencing quality can differ between subtypes within a single bacterial species. While results from this study may not be broadly generalisable, it underscores that a detailed understanding of the driving mechanisms behind ONT sequencing and assembly errors is essential to predict performance and guide methodological choices in future genomic studies.

## Acknowledgements

We want to thank employees at the Department of Medical Microbiology, St Olavs Hospital, for supporting this project. We also want to thank clinicians and other employees at Nord-Trøndelag Hospital Trust for their support and contribution to data collection.

## Supporting information

Supplementary Table S1. Number of concordant and discordant presence and total copy number of genes in ONT and Illumina assemblies compared to polished ONT assemblies, showing reference genes with GC content, bp length and accession numbers.

Supplementary Table S2. Quality markers of reads and assemblies from isolates belonging to ST25.

Supplementary Table S3. Mapping of selected isolates to reference genes.

Supplementary Table S4. Replicons on contigs containing *ermC* and *blaZ* genes.

Supplementary Figure F1. Streme report.

